# Single-cell geometry of drug responses across malignant glioma states

**DOI:** 10.64898/2026.11.03.763852

**Authors:** Evgeniia Lavrenteva, Kim Kyung Do

## Abstract

Malignant gliomas exhibit substantial transcriptional heterogeneity: AC-, OPC-, NPC- and MES-like states can coexist within the same tumor, and a drug affecting one state may leave others relatively unaffected. Whether these states respond in a shared direction, or diverge, is rarely tested directly, since drug response is usually summarized by one score that collapses this heterogeneity. A public single-cell RNA-sequencing resource of drug-treated glioma slice cultures (391,444 cells) is reanalyzed here to test this directly. After restricting to 252,106 transformed glioma cells, 43 treatment-vehicle comparisons from 19 TissueID-defined units are evaluated for response-direction coherence (C, similarity of state-conditioned treatment vectors) and change in state separation (K, state-centroid contraction or expansion). C proves to be the reproducible signal: it survives high-confidence state filtering (r=0.931), an independently selected 500-gene space (r=0.634), the published Neftel modules (r=0.783) and non-overlapping gene cross-fitting (r=0.809), and exceeds two null models in 41 of 43 comparisons, a result that also holds across the 19 biological units themselves (19 of 19 positive; P=3.8e-6). Within-drug coherence exceeds between-drug coherence in all 52 drug pairs and all 11 multi-drug units (P=0.000488): response direction is drug-specific, not generic. K tells a different story: stable only when the feature space is fixed, it loses concordance whenever that space shifts, with only about half of comparisons separating from the null. Malignant glioma states largely agree on the direction of drug response even when they disagree on its magnitude.

## Introduction

Glioblastoma is the most common and most lethal primary malignant brain tumor in adults. The current standard of care, concurrent radiotherapy and temozolomide followed by adjuvant temozolomide, extends median survival from 12.1 to only 14.6 months relative to radiotherapy alone (Stupp et al., 2005), and recurrence after this regimen is effectively universal. A growing body of work attributes this resistance less to additional genetic evolution than to the tumor’s capacity for phenotypic plasticity, in which malignant cells shift reversibly among transcriptional states under treatment and microenvironmental pressure, allowing the tumor to persist even when a dominant state is effectively targeted (Yabo et al., 2022). Quantifying drug response in glioblastoma therefore has to contend with this state-level heterogeneity directly, rather than treating the tumor as a single, uniform population.

Bulk transcriptomic studies established recurrent glioblastoma expression subtypes and showed that subtype composition can change during tumor evolution (Verhaak et al., 2010; Wang et al., 2017). Single-cell studies then showed that these labels do not define mutually exclusive malignant populations: individual tumors contain mixtures of transcriptional programs, including AC-like, OPC-like, NPC-like and MES-like states, together with hybrid and intermediate cells (Patel et al., 2014; Neftel et al., 2019). Subsequent work connected this plasticity to developmental hierarchies, injury-response programs, mesenchymal transitions, stem-like programs, spatial organization, invasion and therapy (Darmanis et al., 2017; Couturier et al., 2020; Yu et al., 2020; Castellan et al., 2021; Hara et al., 2021; Richards et al., 2021; Ravi et al., 2022; Wang et al., 2022; Eisenbarth and Wang, 2023; Pérez-González et al., 2023; Doroszko et al., 2025). This makes drug response a state-resolved problem: the same perturbation can act on several malignant programs that coexist within one biological unit.

Acute patient-derived glioma slices preserve aspects of local tissue architecture while permitting ex vivo perturbation. Zhao et al. used this system with single-cell RNA sequencing to resolve cell-type-specific drug responses (Zhao et al., 2021). Levitin et al. subsequently integrated these and additional experiments using consensus scHPF (Levitin et al., 2019; Levitin et al., 2023), producing a resource spanning 19 reported patients, 10 treatment conditions, 52 samples and approximately 400,000 cells.

Existing computational approaches to single-cell perturbation data address a related but distinct set of questions. scGen, CellOT, CPA and Squidiff model or predict treated expression distributions from unperturbed profiles; scPerturb and scArchon provide harmonized data and benchmarking resources for such models; and scFOCAL links glioblastoma state expression to drug signatures (Lotfollahi et al., 2019; Bunne et al., 2023; Lotfollahi et al., 2023; Peidli et al., 2024; Ahlmann-Eltze and Huber, 2025; He et al., 2026; Radig et al., 2026; Suter et al., 2026). These methods are built to predict or characterize an expression response, not to ask whether several coexisting malignant programs move through transcriptional space in a shared direction once a response is already measured. More generally, single-cell data science has emphasized that forcing continuous biological variation into discrete categories can affect downstream interpretation (Lähnemann et al., 2020). The question addressed here is narrower: when several recurrent malignant programs are present in the same biological unit, do their transcriptional responses point in similar directions, and does treatment make those programs more or less separated from one another?

The aim is therefore different from perturbation prediction, differential-expression testing, or drug-efficacy ranking. Rather than predicting an unobserved treated profile, responses that are directly measured in matched treated and vehicle samples are summarized, asking how those responses relate across recurrent malignant programs. The three quantities are deliberately kept separate: C captures directional agreement, K captures change in state-centroid separation in a stated feature space, and M captures response magnitude. This separation is intended to avoid treating a large response, a coordinated response, and a change in state geometry as the same phenomenon.

Response direction and state separation are therefore treated as two different quantities rather than combined into one score. Response-direction coherence (C) measures the similarity of state-conditioned treatment vectors. Change in state separation (K) measures the treatment-associated change in the distances among state centroids, while mean response magnitude (M) is retained separately. Whether the conclusions survive changes in state confidence and representation is then assessed: high-confidence state filtering, an independently selected 500-gene space, the published Neftel modules and non-overlapping gene cross-fitting. These checks establish relative C as the main comparative result and K as a feature-space-specific quantity.

## Materials and Methods

### Dataset and definition of biological units

The integrated glioma single-cell RNA-sequencing loom released with the consensus scHPF study (Levitin et al., 2023) is analyzed; it contains drug-perturbed human glioma slice cultures from 19 reported patients, 10 treatment conditions, and 52 samples, including data from the previously published GSE148842 dataset (Zhao et al., 2021). The analysis is restricted to cells annotated by the source study as transformed glioma cells; no new tissue was collected, and malignant status was not independently reassigned. The loom provides Sequencing_Name, Treatment, Cell_type, and TissueID metadata. Although Sequencing_Name contains 22 PW-prefixed identifiers, these correspond to 19 distinct TissueID values, with several PW identifiers mapping to the same TissueID and PW036 containing two TissueIDs. Because the 19 TissueIDs match the 19 biological units reported in the source study, TissueID was used as the biological unit for within-unit standardization, feature selection, geometric analyses, and across-unit summaries. PW identifiers were retained only for provenance and visualization. Since no additional patient-level identifier is available in the loom, TissueID-defined biological units are referred to throughout rather than independently verified individual patients.

### Normalization

No per-cell library-size field is present in the loom. Total counts were calculated from all 60,725 genes, cached, scaled to 10,000 molecules per cell and transformed as log(1 + scaled count), following standard single-cell RNA-sequencing practice (Lun et al., 2016). For cell i and gene g:

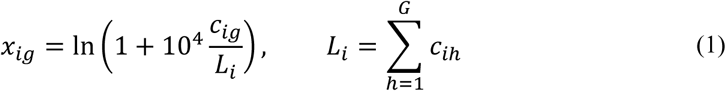

where c_ig_ is the raw count, L_i_ is the total count for cell i, and G = 60,725. Library size was calculated from the full matrix, independently of the marker panel used for state geometry.

### Compact malignant-state scores

For the primary analysis, a compact 34-gene representation spanning AC-like, OPC-like, NPC-like and MES-like programs (7, 7, 10 and 10 genes, respectively; Supplementary Table S2) is assembled. The panel draws on canonical features reported across glioblastoma studies (Neftel et al., 2019; Couturier et al., 2020; Castellan et al., 2021; Hara et al., 2021; Richards et al., 2021; Ravi et al., 2022; Wang et al., 2022; Doroszko et al., 2025) and was used as a deliberately small, interpretable coordinate system for all primary comparisons. It was not varied by drug or TissueID, and all 34 genes were detected in the loom. Because any compact marker panel can impose its own geometry, the analysis is also repeated with the broader published Neftel modules and with an independent 500-gene feature space.

### State assignment

For each TissueID and marker gene, normalized expression was standardized against vehicle-treated transformed cells from the same biological unit. The vehicle mean and standard deviation are denoted in Eq. (2):

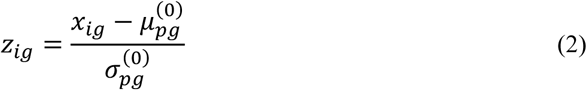

A near-zero or non-finite vehicle standard deviation was replaced by 1. Each state score was the mean standardized expression across its marker set, and the highest-scoring program was assigned to the cell:

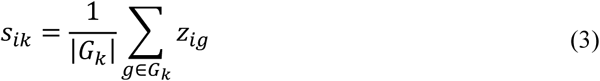

Vehicle and treated cells were assigned independently. These hard labels are an operational stratification of a continuous malignant-state landscape, not a claim that the four programs are mutually exclusive cell types. They define state-conditioned population contrasts; they do not track the same cells through time or estimate transition probabilities. Dependence on this operational labeling was evaluated by high-confidence filtering, the broader published Neftel modules, and non-overlapping cross-fitting described below.

### Inclusion criteria

A TissueID entered paired analysis with at least 100 transformed vehicle cells; a treated condition required at least 50 cells. A state was retained with at least 25 cells in both conditions, and at least three states were required for geometry. These rules were applied uniformly across TissueIDs and drugs. No comparison was removed because of its C, K, M or visual extremity. Forty-three comparisons met the criteria: 41 with four retained states and two with three.

### State-specific treatment vectors

For each retained state, biological unit and condition, the centroid was the mean standardized expression vector over all cells assigned to that state and condition:

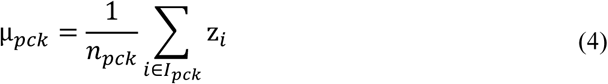

For drug d, the state-conditioned treatment vector was the difference between the treated and matched vehicle centroids:

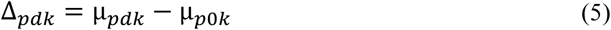

The same definitions were used in all sensitivity analyses; only the genes defining the coordinates and, where specified, the state labels changed.

### Response-direction coherence (C)

For a comparison containing m retained states, response-direction coherence (C) was defined as the mean pairwise cosine similarity among state-conditioned treatment vectors. Cosine similarity was chosen because the question is whether state responses point in a common direction independently of their length. A common positive rescaling of either vector therefore does not change the pairwise value. In contrast, Euclidean distance would mix direction with response magnitude, and Pearson correlation would first center the gene coordinates and would therefore answer a different question about relative gene-wise pattern rather than the angle between the treated-minus-vehicle vectors themselves. For each state pair, cosine similarity was calculated only when the product of the Euclidean norms exceeded epsilon = 10⁻⁸:

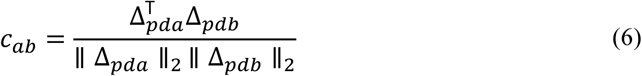

Coherence was then the arithmetic mean of the valid pairwise cosine similarities:

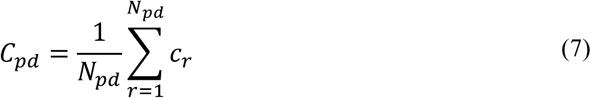

Epsilon was used only to determine whether a denominator was sufficiently non-zero and was not added to the vector norms. C approaches +1 for parallel responses, is near 0 for weak directional agreement and can be negative for opposing responses. A common positive rescaling leaves C unchanged, although changing the biological feature space can alter it.

### Change in state separation (K)

For the same m retained states, mean pairwise Euclidean distance among the state centroids was calculated separately for vehicle (c=0) and treatment (c=d):

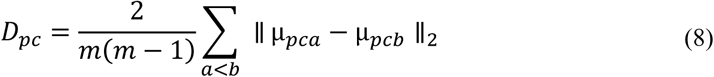

Change in state separation was expressed as a base-2 log-ratio:

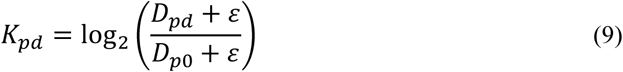

Here, epsilon *ε* = 10⁻⁸ is the numerical safeguard used in the MATLAB analysis. K < 0 indicates reduced state-centroid separation after treatment, K = 0 no change and K > 0 greater separation in the specified feature space. Because Euclidean distance depends on the coordinates, K was treated as representation-specific.

### Response magnitude and state proportions

Mean response magnitude was calculated independently of C and K as the average Euclidean length of the retained state-response vectors:

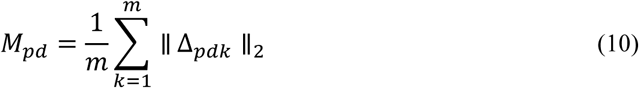

State composition was summarized secondarily by the total variation distance between the treated and vehicle state-proportion vectors (Gibbs and Su, 2002):

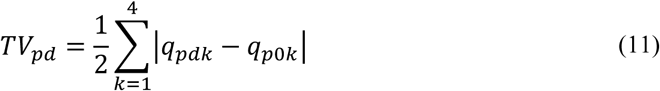

where q_pdk_ and q_p0k_ are the treated and vehicle proportions assigned to state k. TV ranges from 0 to 1 and was used only as a secondary descriptor; it does not enter C, K or M.

### State filtering

As a sensitivity analysis, only cells whose largest compact module score exceeded the second-largest by at least 0.25 are retained. This asks whether the primary result depends mainly on cells lying close to a hard state-assignment boundary. The same inclusion rules and geometry definitions were then reapplied. Concordance with the primary analysis was summarized by Pearson correlation, sign agreement, median absolute change and median signed change.

### Independent 500-gene feature space

A second sensitivity analysis changed the coordinates used to measure response while keeping the compact state labels fixed. Highly variable genes were selected from vehicle-treated transformed cells only. For bounded HDF5 access, up to two contiguous vehicle-cell blocks were sampled per TissueID, with up to 75 cells from each block (maximum 150 per unit; 1,950 cells in total). The 34 compact markers, mitochondrial genes, ribosomal protein genes and MALAT1 were excluded. Genes detected in at least 2% of sampled cells were ranked by average within-TissueID variance, and the top 500 were used to recalculate C, K and M. The 500-gene size and 2% detection threshold were used uniformly in this analysis. This is a within-cohort feature-space sensitivity analysis, not an independent biological validation.

### Published Neftel modules

As a broader alternative to the 34-gene panel, the published Neftel meta-module gene sets (Neftel et al., 2019) are used. Detected module genes were scored with the same within-TissueID vehicle-referenced standardization used for the compact analysis. NPC1 and NPC2 were combined as NPC-like, and MES1 and MES2 as MES-like, yielding the same four broad state classes for comparison with the primary representation. C, K and M were then recalculated in this broader representation. This analysis asks whether the main result depends on the compact marker definition rather than treating the Neftel labels as an independent biological validation.

### Non-overlapping cross-fit

To test a more specific source of circularity, gene overlap between state assignment and response geometry is removed. Detected Neftel genes were partitioned globally into disjoint sets A and B in ten deterministic pseudo-random splits (base seed 19062019), with at least 12 genes from every meta-module in each half. Global partitioning ensured that a gene shared by modules could not enter assignment through one membership and geometry through another. The 12-gene minimum was a design constraint to keep every meta-module represented, not a biological cutoff. Each split was evaluated reciprocally: A defined state labels while B defined geometry, then the roles were reversed, producing 20 held-out folds. For each biological-unit-drug pair, the median across valid folds was retained, with quartiles as a stability summary. A pair entered the final comparison when the metric was calculable in at least 75% of folds. Assignment stability was summarized by raw agreement and Cohen’s kappa (Cohen, 1960).

### Within-unit null-model calibration

Because C, K and M are descriptive geometric quantities, their absolute values were calibrated with two within-unit null models constructed separately for each biological-unit-drug comparison. In the vehicle-split null, vehicle cells within each retained state were randomly partitioned into pseudo-vehicle and pseudo-treated groups matched to the observed treated-to-vehicle cell-count ratio. In the label-permutation null, treatment labels were shuffled within state while reproducing the observed per-state group sizes. Each null model used 1,000 randomizations per comparison. C, K and M were recalculated under every randomization. Empirical P values for C and M were calculated as (1 + number of null statistics greater than or equal to the observed statistic)/(B + 1), with B = 1,000; departure of K from zero was evaluated in absolute value because either contraction or expansion can produce a large deviation. These comparison-level P values were used as calibration summaries rather than as a multiple-testing framework for declaring individual drug effects. The nulls assess whether the observed geometry exceeds sampling variation at the available cell numbers; they do not constitute independent biological replication or reproduce slice-to-slice biological variability.

### Cross-drug specificity analysis

For the specificity analysis, biological units contributing at least two drug conditions were considered. For each within-unit drug pair, within-drug C was compared with cosine similarity between state-conditioned response vectors from different drugs. Same-state between-drug similarity is also calculated as a complementary comparison. Because multiple drug pairs can arise from the same TissueID, the 52 pairwise comparisons are nested within 11 biological units. Pair-level values were therefore treated as descriptive contrasts rather than independent biological replicates. To obtain a biological-unit-level summary, the median difference between within-drug coherence and between-drug coherence across its available drug pairs is calculated within each TissueID. These 11 TissueID-level differences were summarized by their median and a 95% percentile bootstrap interval from 2,000 resamples of biological units. A one-sided exact sign test evaluated whether the TissueID-level differences were greater than zero, matching the directional specificity contrast of within-drug versus between-drug coherence. The same procedure was applied to the complementary comparison using same-state between-drug similarity. These analyses distinguish coherence shared by states under one perturbation from a generic response direction shared across different perturbations; they were not used to rank drugs.

Two further checks addressed alternative explanations for C. To test whether the observed coherence could reflect a generic dissociation-stress response rather than a drug-specific effect, a per-cell dissociation-stress score was computed from 11 canonical stress-response genes (FOS, FOSB, JUN, JUNB, EGR1, HSPA1A, HSPA1B, ZFP36, IER2, NR4A1, DUSP1; van den Brink et al., 2017), each gene standardized against its patient-matched vehicle population exactly as in Eq. (2), and the mean z-score across the panel averaged per treated population; this score was correlated with C across all 43 comparisons using Pearson’s r. To test whether between-drug coherence was specific to shared mechanism of action rather than a general perturbation axis, each within-unit drug pair from the cross-drug analysis above was labeled by whether its two drugs shared a mechanism-of-action class (HDAC inhibitor, topoisomerase inhibitor, GPX4 inhibitor/ferroptosis inducer, TrkB antagonist, gamma-secretase inhibitor, or KSP/Eg5 inhibitor, assigned from the drugs’ published mechanisms), and between-drug coherence was compared between same-mechanism and different-mechanism pairs with a two-sided Wilcoxon rank-sum test.

### Drug-level summaries and statistics

Cells were not treated as independent biological replicates. Across-unit drug summaries used TissueID as the biological unit and were reported only for drugs represented in at least three TissueIDs. Medians were accompanied by 95% percentile bootstrap intervals from 2,000 resamples of biological units (Efron, 1979). Because each drug was represented by only 3-8 units, these intervals are used as limited-sample uncertainty summaries.

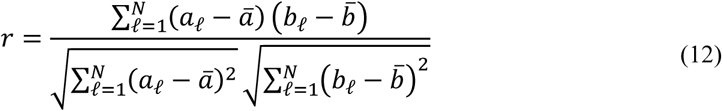

Pearson correlation was used descriptively to compare primary and sensitivity-analysis metrics across biological-unit-drug comparisons. Because some TissueIDs contribute more than one drug condition, no independence-based P values were attached to these correlations. For within-TissueID centered correlations, the TissueID-specific mean of each metric was subtracted from every observation for that TissueID before the pooled Pearson correlation was calculated. For K, sign consistency relative to zero was assessed with the exact two-sided sign test (Dixon and Mood, 1946). If B is the number of positive non-zero K values among n units:

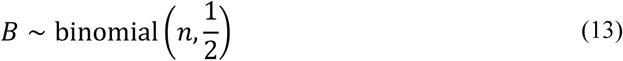

The two-sided P value was obtained from the corresponding binomial tail, with zeros omitted. Assignment stability was described by raw agreement and Cohen’s kappa (Cohen, 1960):

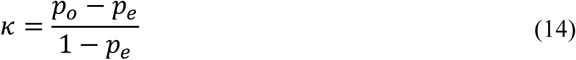

where observed and chance-expected agreement are defined in Eq. (14). All reported comparisons use the same prespecified inclusion rules stated above, and no metric-based outlier exclusion was performed. Drug-level summaries and cross-analysis correlations are interpreted descriptively because of the limited number of biological units and repeated drug conditions within some TissueIDs.

## Results

### Dataset and biological units

The integrated loom contained 391,444 cells, of which 252,106 were annotated as transformed glioma cells with usable treatment metadata and a matched vehicle reference. Identifier auditing showed 22 PW prefixes but exactly 19 TissueID values. Several PW prefixes mapped to the same TissueID, and PW036 contained two TissueIDs; TissueID was therefore used as the biological grouping variable. PW030, used below as the multi-drug example, mapped uniquely to TB5884. Applying the inclusion criteria yielded 43 matched treatment-vehicle comparisons: 41 retained all four canonical states and two retained three. Nine drugs were represented in at least three TissueID-defined units: ANA-12, RO4929097, etoposide, givinostat, ispinesib, panobinostat, RSL3, RSL3 plus ferrostatin, and topotecan. Large cell numbers improve within-unit centroid estimation but do not increase the number of biological replicates.

### Definition of C, K and M

Within each TissueID, treatment vectors were defined as treated-minus-vehicle state centroids. C summarizes pairwise similarity among their directions, K the log2 treated-to-vehicle change in mean state-centroid separation, and M their mean magnitude. Because state assignment was recomputed after treatment, these are state-conditioned population contrasts rather than cell trajectories. Across 43 comparisons, C ranged from 0.059 to 0.856 (median 0.608) and was positive throughout. K ranged from −0.260 to 1.092 (median 0.044), with 28 positive and 15 negative values (Supplementary Figure S2; Supplementary Table S1). Relative robustness was assessed by changing state definitions and expression coordinates; absolute C, K and M were additionally calibrated against the within-unit null models reported below.

### Drug responses in TB5884 (PW030)

TB5884 (PW030) contained matched vehicle slices and five drug conditions under the same within-unit standardization. ANA-12, RO4929097, etoposide and ispinesib showed C = 0.736-0.856 and K = 0.113-0.186. Panobinostat differed from the other four conditions, with C = 0.624, K = 0.613 and the largest response magnitude (M = 2.583). Its K value corresponds to a treated-to-vehicle centroid-distance ratio of 1.529, or approximately 53% greater mean pairwise separation in the compact 34-gene representation (Figure 1). This unit is shown because it illustrates, within one biological unit, that C, K and M need not move together: the largest response magnitude and state-space expansion did not coincide with the highest directional coherence. It is not representative of the typical panobinostat response: among six units exposed to panobinostat, median K was 0.010 (95% bootstrap interval −0.110 to 0.372), with three positive and three negative values, and median C was 0.606, close to the all-drug median. The pronounced K and M seen in TB5884 are therefore a property of this unit rather than a consistent panobinostat effect (Figure 3; Table 1).

**Figure 1.**
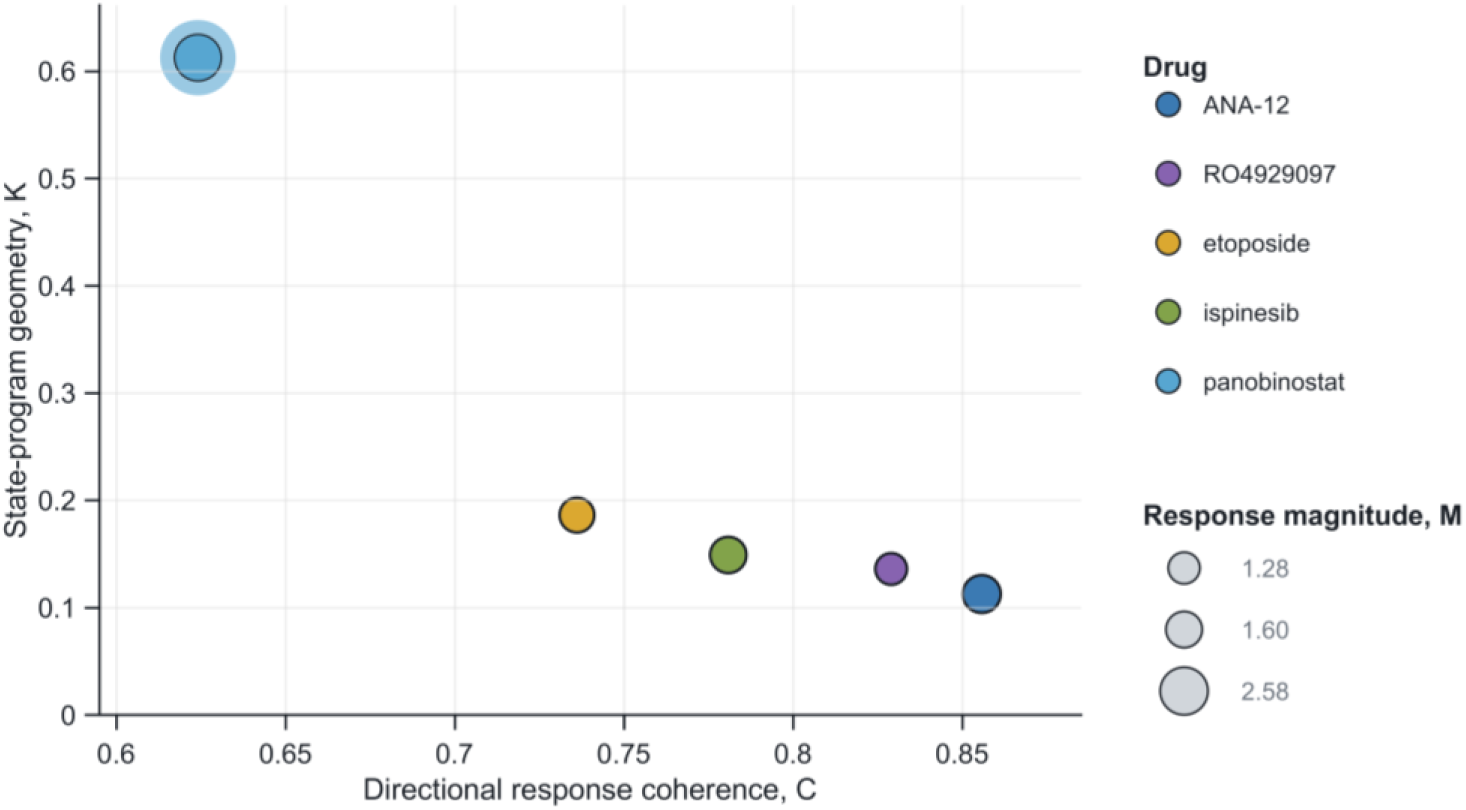
Each point is one drug condition relative to the TissueID-matched vehicle reference. C is response-direction coherence, K is the log2 treated-to-vehicle ratio of mean pairwise state-centroid distance, and point size represents mean response magnitude M. Positive K indicates greater and negative K reduced separation in the compact 34-gene representation.

**Table 1.** Biological-unit summary for drugs represented in at least three TissueID-defined units.

| **Drug** | **n** | **Median C** | **C 95% bootstrap interval** | **Median K** | **K 95% bootstrap interval** | **K sign consistency** | **Exact sign P** | **Median M** |
|---|---|---|---|---|---|---|---|---|
| ANA-12 | 3 | 0.771 | 0.730 to 0.856 | 0.113 | 0.040 to 0.137 | 1.000 | 0.250 | 1.211 |
| RO4929097 | 3 | 0.682 | 0.595 to 0.829 | 0.005 | -0.081 to 0.136 | 0.667 | 1.000 | 1.258 |
| Etoposide | 6 | 0.674 | 0.390 to 0.764 | 0.034 | -0.137 to 0.135 | 0.667 | 0.688 | 1.020 |
| Givinostat | 4 | 0.533 | 0.059 to 0.735 | 0.067 | 0.038 to 0.166 | 1.000 | 0.125 | 0.789 |
| Ispinesib | 5 | 0.779 | 0.255 to 0.808 | -0.085 | -0.207 to 0.149 | 0.600 | 1.000 | 1.175 |
| Panobinostat | 6 | 0.606 | 0.478 to 0.650 | 0.010 | -0.110 to 0.372 | 0.500 | 1.000 | 1.235 |
| RSL3 | 8 | 0.443 | 0.371 to 0.564 | 0.056 | -0.018 to 0.206 | 0.625 | 0.727 | 0.945 |
| RSL3 + ferrostatin | 3 | 0.264 | 0.155 to 0.458 | 0.113 | -0.033 to 0.200 | 0.667 | 1.000 | 0.696 |
| Topotecan | 5 | 0.636 | 0.544 to 0.723 | 0.150 | -0.009 to 0.266 | 0.600 | 1.000 | 1.385 |

Bootstrap intervals are percentile intervals from 2,000 resamples of biological units. The exact two-sided sign test is applied only to K relative to zero. With 3-8 biological units per drug, intervals and P values summarize cross-unit consistency rather than support drug ranking. For the two drugs represented by only three units (ANA-12, RO4929097), resampling from three observations admits few distinct bootstrap compositions, so the reported intervals for those rows should be read as coarse, approximate ranges rather than well-calibrated confidence intervals. Null-model randomization and empirical-P calculations are described in Materials and Methods.

### Response-direction coherence by drug

All nine replicated drugs had positive median C, although the magnitude varied substantially (Figure 2). Median C ranged from 0.264 for RSL3 plus ferrostatin to 0.779 for ispinesib; ANA-12 was similarly high at 0.771. These values describe a shared directional component of the transcriptional response and do not imply identical state responses, therapeutic efficacy or significant differences between drugs.

**Figure 2.**
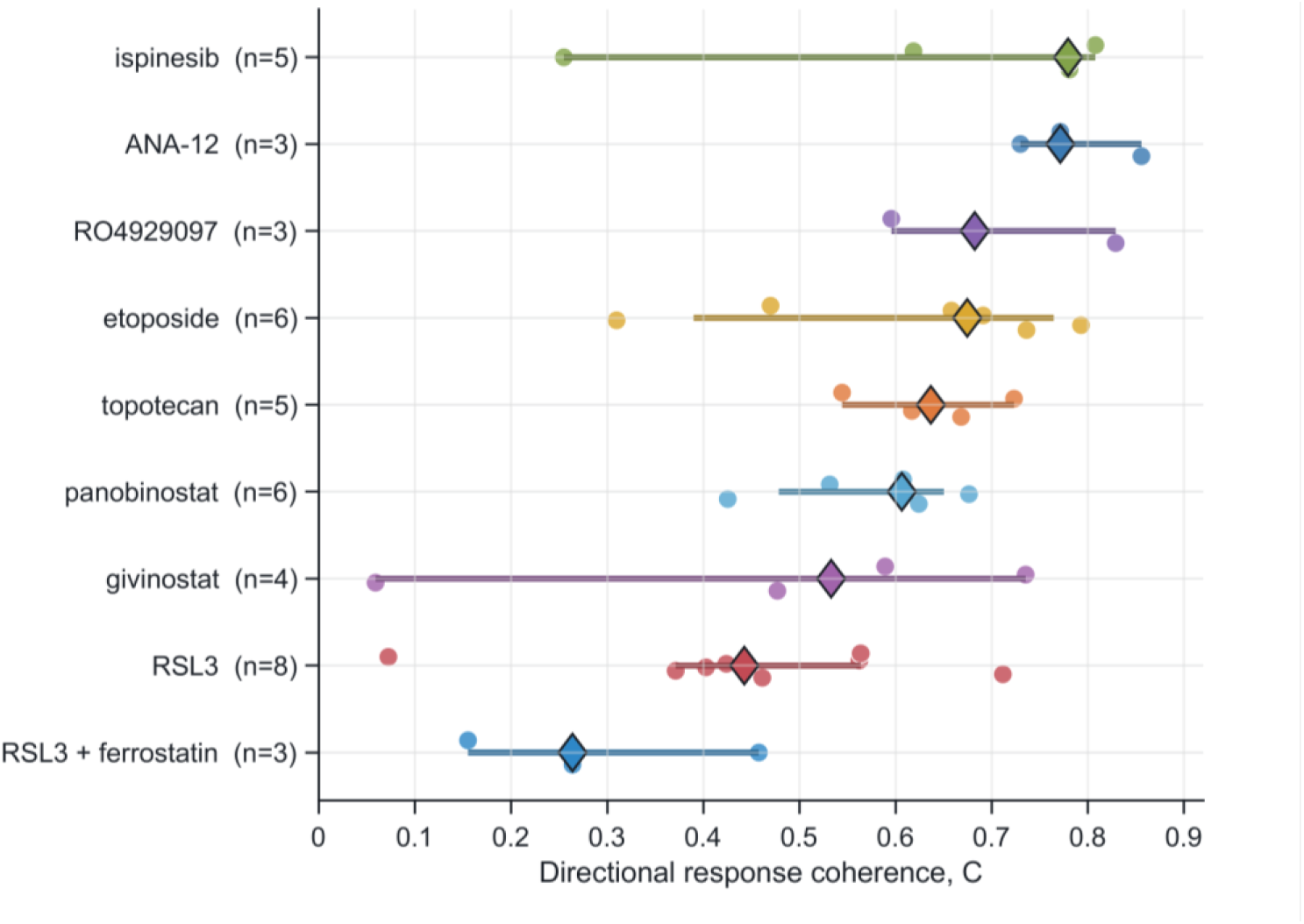
Response-direction coherence for the nine drugs represented in at least three TissueID-defined biological units. Circles show individual biological-unit estimates, diamonds show medians, and horizontal intervals are 95% percentile bootstrap intervals from 2,000 resamples of biological units.

### Change in state separation by drug

K showed much weaker cross-unit consistency (Figure 3; Table 1). Median K ranged from −0.085 for ispinesib to 0.150 for topotecan, and most bootstrap intervals included zero. ANA-12 and givinostat had positive K in all available units, but these estimates were based on only three and four units, respectively. Panobinostat had three positive and three negative values. None of the replicated drugs showed convincing evidence of consistent contraction or expansion across biological units. Across the 43 comparisons, K correlated strongly with response magnitude M (Pearson r=0.756; r=0.833 after centering within biological unit) and moderately with the total variation distance between treated and vehicle state proportions defined above (r=0.489; r=0.478 within-unit), so that comparisons with a larger overall response also tended to show more state-composition change and more state-centroid separation. Because K and M are both derived from Euclidean displacements in the same representation, some association between them is mathematically expected; these correlations are therefore descriptive rather than evidence that response magnitude causally determines K. The result nevertheless shows that K is not independent of overall perturbation magnitude, which is distinct from the directional coherence question addressed by C. After removing the linear dependence of K on M by regression, the residual K retained 43% of its original variance, and this residual did not correlate with the state-composition shift either (r=-0.070, P=0.656). K therefore is not simply a rescaled copy of M, but given K’s already weak cross-unit reproducibility (above), this residual variance is better characterized as unexplained instability than as a confirmed independent signal.

**Figure 3.**
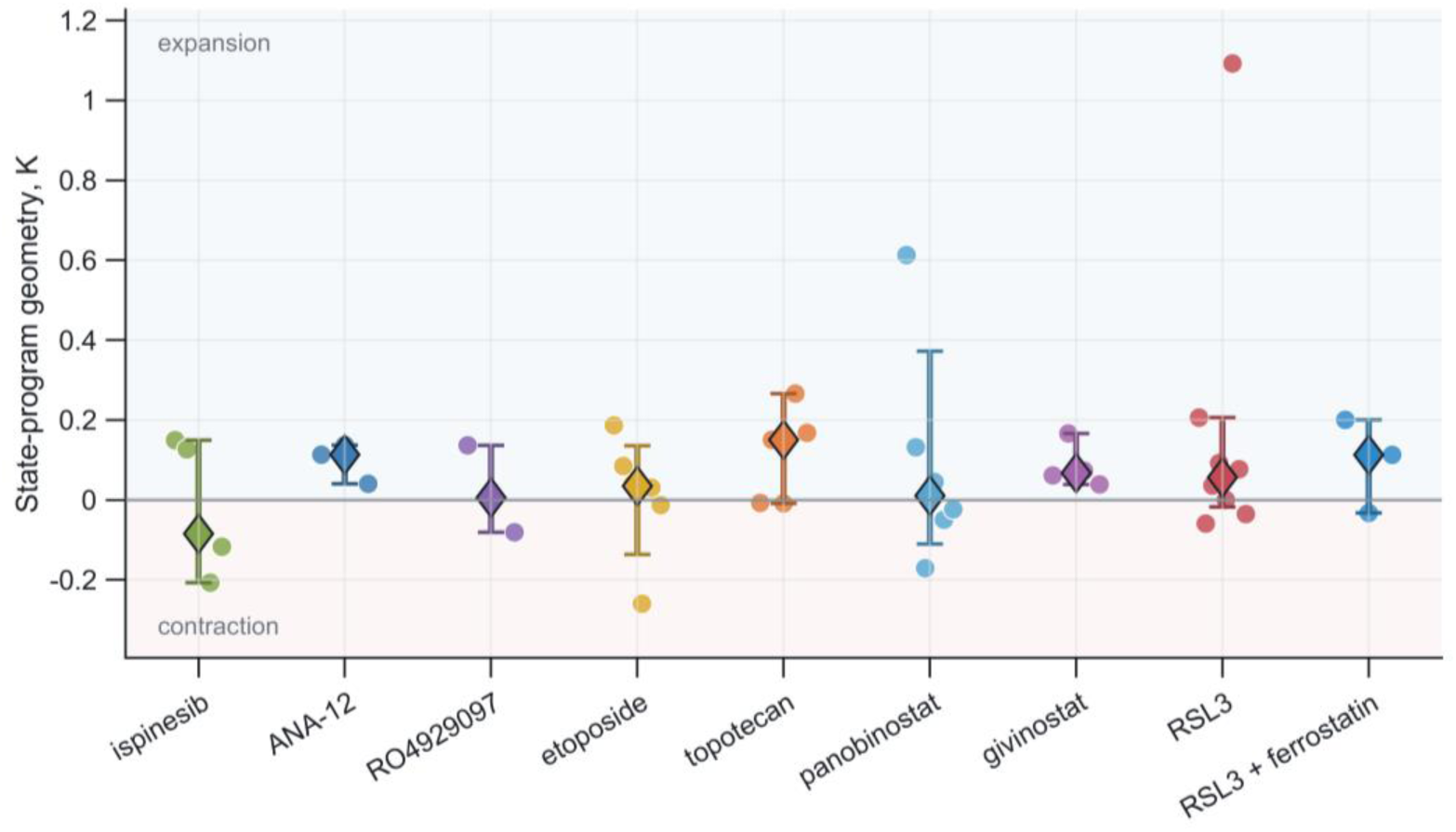
Change in state separation K for the same nine drugs. Circles show individual biological-unit estimates, diamonds show medians, and vertical intervals are 95% percentile bootstrap intervals. The zero line denotes no change; positive K indicates greater separation and negative K reduced separation in the compact representation.

### High-confidence state filtering

After retaining only high-confidence compact-state assignments, 42 comparisons remained evaluable. C was strongly concordant with the primary analysis (r=0.931), with no sign reversals and a median absolute change of 0.168; absolute C values shifted downward (median signed change −0.168). This downward shift is consistent with boundary cells — those with similar scores for two or more states — contributing a shared, less state-specific component to the standardized expression profile; removing them should sharpen state-specific signal and so reduce the common directional component that enters C, without reversing its sign. K was also stable within the unchanged compact feature space (r=0.918; 88.1% sign agreement; median absolute change 0.048). The primary pattern was therefore not driven mainly by cells with near-tied state scores (Figure 4; Supplementary Table S3).

**Figure 4.**
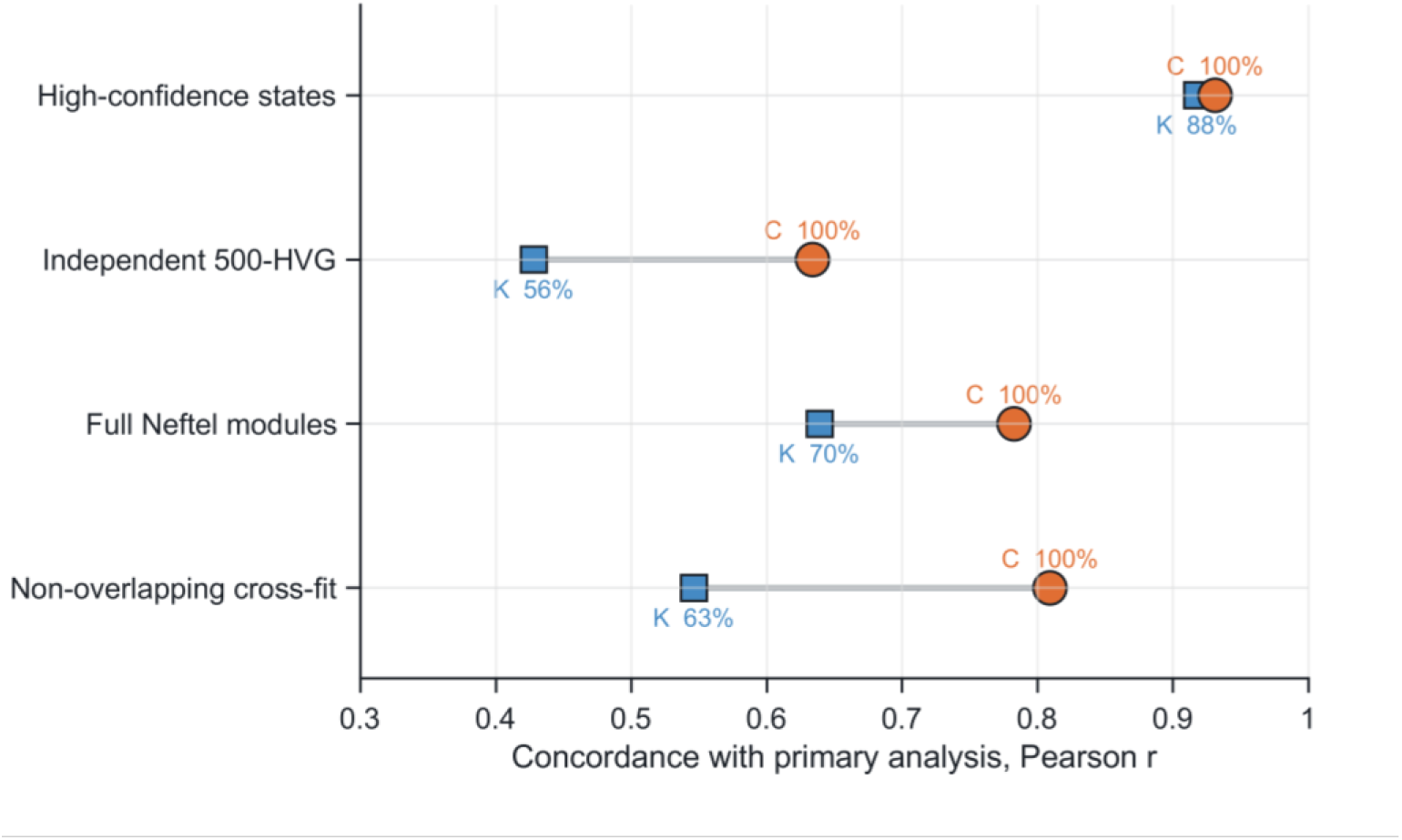
Concordance of C and K between the primary analysis and four sensitivity analyses. Horizontal position is the Pearson correlation with primary biological-unit-drug values; percentages show sign agreement. High-confidence filtering retains the compact feature space, whereas the 500-HVG, full Neftel-module and cross-fit analyses change the representation.

### Independent 500-gene feature space

To separate state assignment from the coordinates used to measure response, geometry is recalculated in 500 highly variable genes selected from vehicle cells only, excluding the compact markers and common technical gene classes. Compact state labels are held fixed. C remained moderately concordant with the primary analysis across all 43 comparisons (r=0.634), with no sign reversals, although its absolute scale shifted upward (median signed difference +0.258). K was much less stable (r=0.428; 55.8% sign agreement; median absolute difference 0.264), without a systematic shift toward contraction or expansion (median signed difference +0.003). The broader feature space retained part of the relative C pattern but not a feature-space-invariant K (Figure 4; Supplementary Figure S1; Supplementary Table S3).

### Published Neftel modules

The compact panel is next replaced with the published Neftel meta-modules (Neftel et al., 2019), with near-complete gene coverage. NPC1/NPC2 and MES1/MES2 were collapsed to four state classes after scoring, providing a broader published representation of the same malignant-state framework. Compact and full-module hard labels agreed for 49.6% of transformed cells (Cohen’s kappa = 0.319; median unit agreement 49.4%), showing how categorical labels can vary when a continuous state landscape is represented by different gene sets (Supplementary Figure S3). The response metrics were more stable: full-module C correlated with primary C (r=0.783) with no sign reversals, whereas K was less concordant (r=0.639; 69.8% sign agreement) (Figure 4; Supplementary Table S3).

### Non-overlapping cross-fit

Repeated two-fold cross-fitting removed overlap between genes used for state assignment and response geometry. Ten global A/B partitions were analyzed in both directions, producing 20 held-out folds; comparisons were retained when evaluable in at least 75% of folds. Hard labels remained unstable across the split (median A/B agreement 45.5%; median kappa 0.230), but cross-fitted C remained strongly associated with primary C across all 43 comparisons (r=0.809), with no sign reversals and a median absolute difference of 0.148. Cross-fitted K was weaker (r=0.546; 62.8% sign agreement; median absolute difference 0.211). Using disjoint genes for assignment and geometry within each fold provides the strongest internal check of gene-set circularity in this study. The same cells are reused, so the cross-fit is not independent biological replication. Across all four sensitivity analyses, C was recovered more consistently than K (Figure 4; Figure 5; Supplementary Table S3).

**Figure 5.**
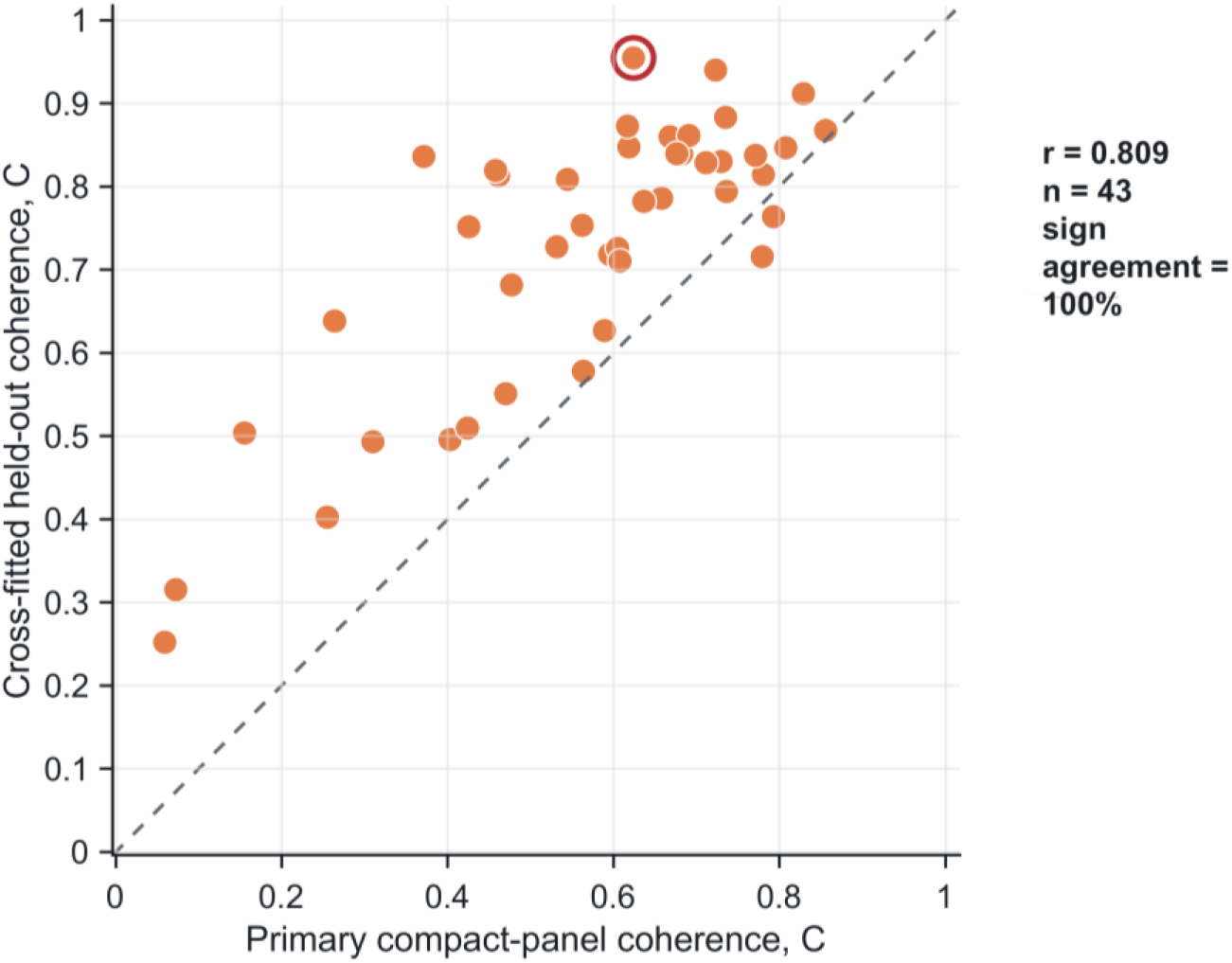
Primary compact-panel C versus median C from reciprocal cross-fits in which genes used for state assignment are disjoint from those used for response geometry. The dashed line is the identity line. Across 43 comparisons, r = 0.809 and no C value changed sign; TB5884-panobinostat is marked to link with Figure 1.

### Calibration of C, K and M against within-unit null models

Because C, K and M are descriptive geometric quantities, their absolute values are calibrated against two null models constructed separately for every biological unit and drug: a vehicle-split null, in which vehicle-treated cells within each retained state were partitioned at random into pseudo-vehicle and pseudo-treated groups matched to the observed cell-count ratio, and a label-permutation null, in which condition labels were shuffled within state while reproducing the observed per-state cell counts exactly. Under the vehicle-split null, median null C was −0.003 against an observed median of 0.608, a median excess of 0.616, and 41 of 43 comparisons had an empirical P below 0.05 (Figure 6). The label-permutation null gave a median null C of −0.004 and also placed 41 of 43 comparisons below 0.05. Observed C therefore exceeds what centroid sampling variation alone produces at these cell numbers. Response magnitude behaved the same way: median M was 1.065 against null medians of 0.407 and 0.343, with all 43 comparisons below 0.05. K did not: only 21 of 43 comparisons under the vehicle-split null and 23 of 43 under the label-permutation null departed from the null in absolute value, consistent with the weak cross-unit consistency reported above. These nulls test only whether the observed geometry exceeds sampling variation; they do not by themselves distinguish a coherent pharmacological response from a single dominant response axis shared across drugs, which is addressed next. The two null models are related constructions built from the same within-unit cell pool — one derives pseudo-replicates from vehicle cells alone, the other permutes condition labels within state — so their close agreement (median null C of −0.003 and −0.004, respectively) reflects consistency between two similar sampling-variation controls rather than two independent lines of evidence for C.

**Figure 6.**
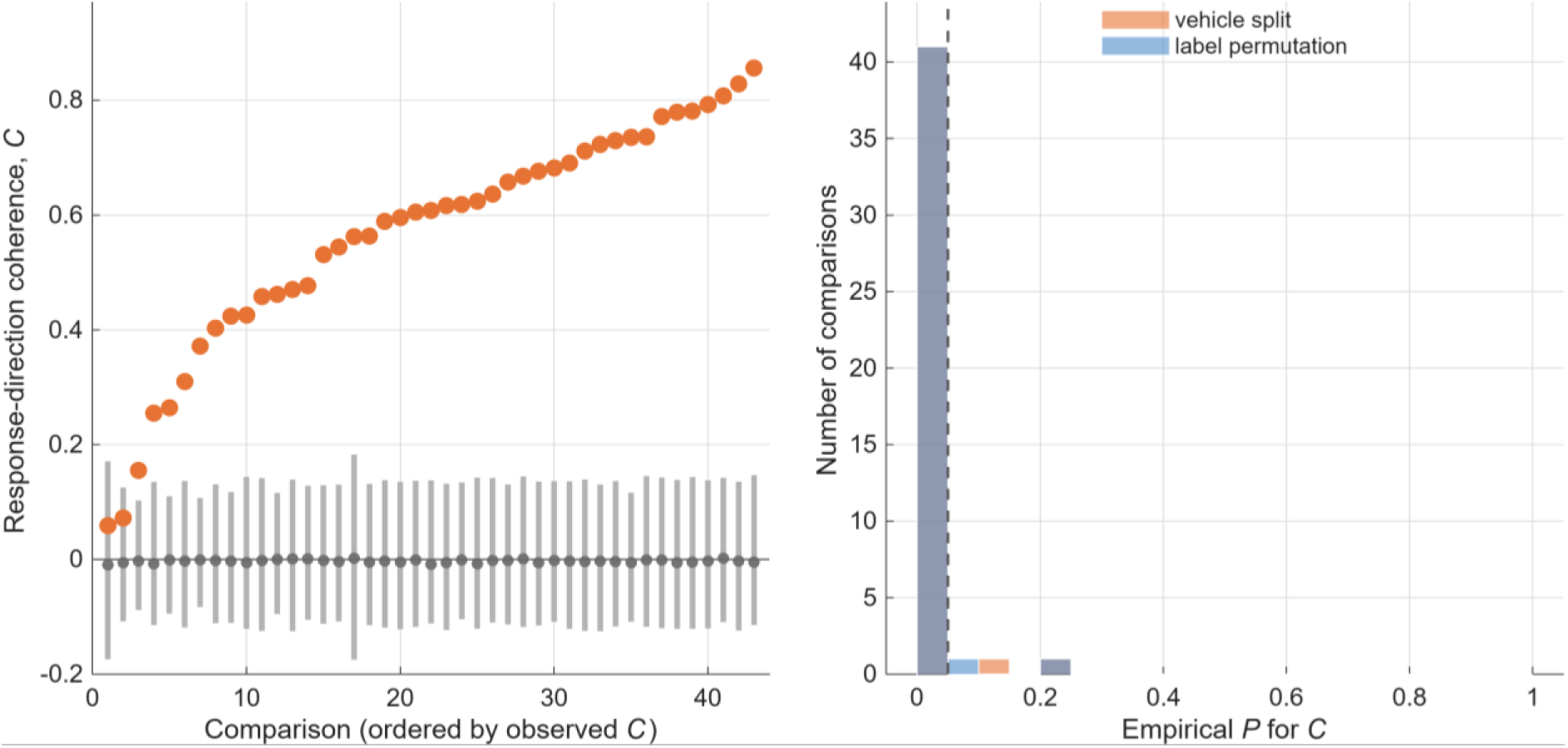
Observed response-direction coherence, C, compared with its vehicle-split null distribution across the 43 comparisons, ordered by observed C (left), and the distribution of empirical P values for C under the vehicle-split and label-permutation nulls across all 43 comparisons (right).

The 43 comparisons are not 43 independent trials: they are nested within 19 biological units, some of which contribute more than one drug comparison. As a unit-level complement to the comparison-level null tests above, one median C was taken per biological unit (collapsing multiple drug comparisons within a unit to a single value) and tested for departure from zero directly, without reference to the null models. All 19 of 19 units had a positive median C (exact two-sided sign-test P=3.8×10-6), and the observed mean unit-level C (0.506, SD 0.212) exceeded the minimum effect this design had 80% power to detect at two-sided alpha=0.05 (0.136) by a factor of 3.7. This unit-level test is independent of the comparison-level null-exceedance count reported above and uses the number of biological units, not the number of comparisons, as its sample size.

### Coherence is largely drug specific

A high C establishes that the retained states move in similar directions under one drug, but not that the shared direction is specific to that drug rather than a general perturbation axis. Across 52 drug pairs in the 11 biological units contributing at least two drugs, median within-drug C was 0.652, whereas the median cosine similarity between response vectors of different states under two different drugs was 0.233, a median pair-level difference of 0.365; within-drug coherence exceeded between-drug coherence in every one of the 52 pairs (Figure 7). Because these pairwise contrasts are nested within TissueIDs, the difference is then summarized once per biological unit. The within-minus-between difference was positive in all 11 TissueIDs, with a median unit-level difference of 0.291 (95% bootstrap CI 0.201-0.495; one-sided exact sign-test P=0.000488). The median same-state between-drug similarity at the pair level was 0.297. In the corresponding unit-level comparison, 9 of 11 TissueIDs had a positive within-minus-between difference, with a median of 0.196 (95% bootstrap CI 0.057-0.430; P=0.0327). Thus, response direction was more strongly associated with perturbation identity than with state identity in these comparisons. The positive between-drug similarities nevertheless indicate a modest response component shared across perturbations.

**Figure 7.**
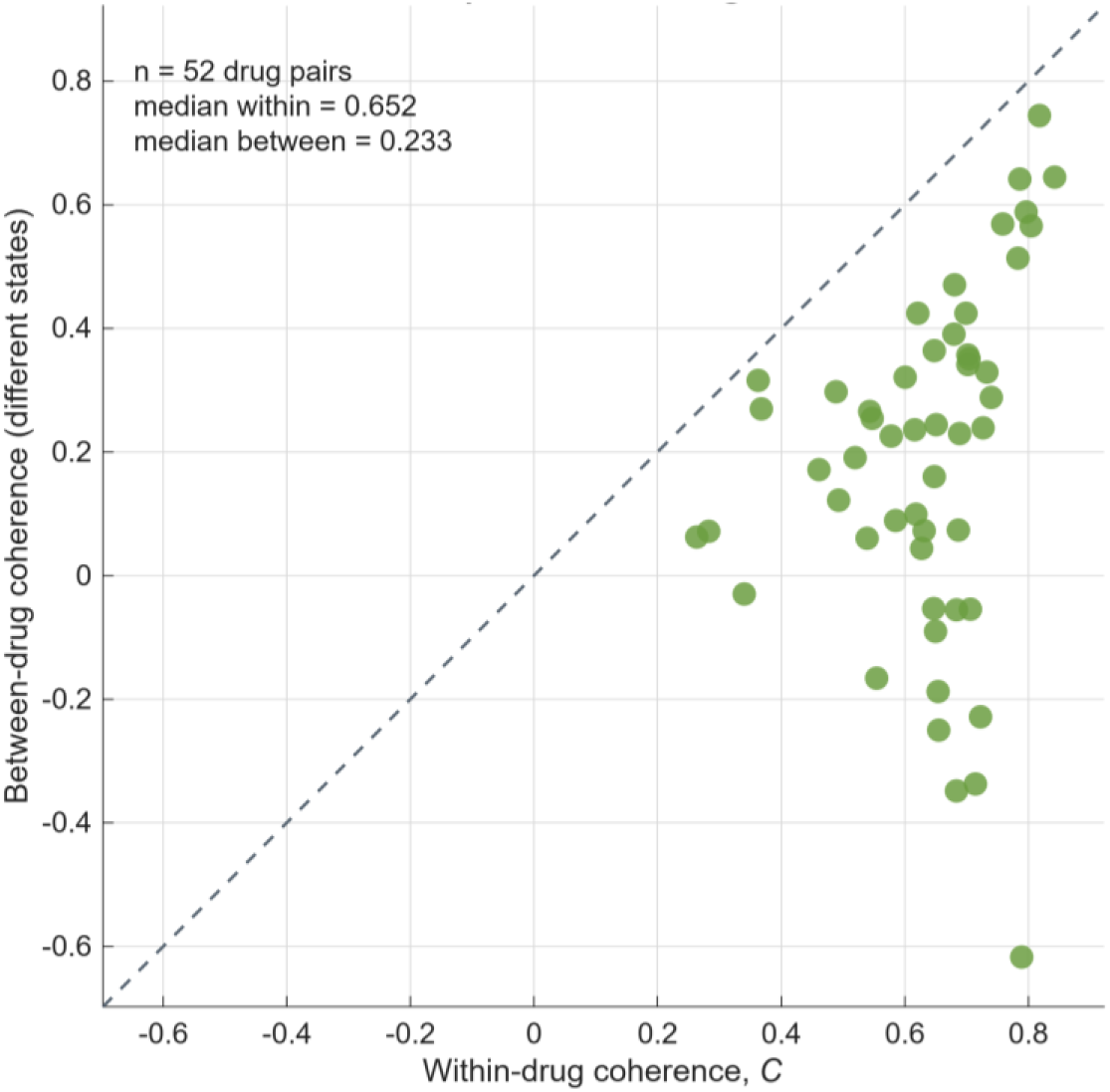
Pair-level comparison of within-drug response-direction coherence with between-drug coherence for different states across 52 drug pairs from 11 biological units.

## Discussion

The clearest result was not that treatment consistently contracted or expanded the malignant-state configuration. It did not. Instead, the relative ordering of response-direction coherence was the part of the analysis that survived changes in how state space was represented. High-confidence filtering gave very similar C estimates, and C remained concordant when geometry was recalculated in 500 independently selected genes, when the full Neftel modules replaced the compact panel, and when assignment genes were separated from geometry genes by cross-fitting. K behaved differently: it was stable when the compact coordinates were retained, but its value and sometimes its sign changed when the coordinate system changed. C is therefore interpreted as the more reproducible comparative summary in this dataset, and K only within an explicitly stated feature space.

This distinction matters because glioblastoma states are recurrent programs rather than fixed, mutually exclusive cell types. Neftel et al. described AC-like, OPC-like, NPC-like and MES-like states while emphasizing hybrid and intermediate cells (Neftel et al., 2019), and later work connected state plasticity to developmental hierarchies, injury responses, mesenchymal transitions, stem-like programs, spatial organization and therapy (Darmanis et al., 2017; Couturier et al., 2020; Yu et al., 2020; Castellan et al., 2021; Hara et al., 2021; Richards et al., 2021; Ravi et al., 2022; Wang et al., 2022; Eisenbarth and Wang, 2023; Pérez-González et al., 2023; Doroszko et al., 2025). In this dataset, compact-panel and full-module hard labels agreed for only about half of transformed cells (49.6%; Cohen’s kappa=0.319, conventionally “fair” agreement). That instability is not surprising for a continuous landscape, and it is one reason the main conclusion is not made to depend on one hard labeling scheme (Lähnemann et al., 2020; Ahlmann-Eltze and Huber, 2025). What was more informative was that relative C persisted despite those changes. C should still be read narrowly: it measures whether state-conditioned treatment vectors share a direction. It does not say that the states respond identically, that the response is beneficial, or that a drug is effective. One unit, TB5884, is used below purely as an illustrative example of this distinction, not as representative evidence for the dataset as a whole. Panobinostat produced the largest response magnitude and a large increase in compact-panel state separation in that unit, but K changed sign across the six biological units exposed to panobinostat. A striking response in one unit was therefore not a reproducible drug-level expansion effect.

The analysis addresses a different question from perturbation prediction. scGen, CellOT, CPA and Squidiff model or predict treated expression distributions, scPerturb and scArchon support harmonization or benchmarking, and scFOCAL links glioblastoma state expression with drug signatures (Lotfollahi et al., 2019; Bunne et al., 2023; Lotfollahi et al., 2023; Peidli et al., 2024; He et al., 2026; Radig et al., 2026; Suter et al., 2026). Here C is not a prediction model, a differential-expression statistic, or an efficacy score; it is an observed-data summary of how measured state-conditioned treatment vectors relate to one another within a biological unit. The cross-fit is important for that interpretation because it asks whether the same genes are manufacturing both the state labels and the apparent response geometry. C remained strongly associated with the primary estimate when assignment and geometry genes were disjoint, whereas K was again less stable. The difference is mathematically expected: cosine direction can remain similar after a change of representation, whereas Euclidean distances are directly tied to the chosen coordinates. For that reason, the large TB5884 panobinostat K value describes expansion of the compact canonical-program representation in that biological unit; it should not be generalized to transcriptome-wide heterogeneity. The present analysis therefore asks whether a coherent response direction exists and is reproducible across representations; it does not decompose that direction into gene- or pathway-specific mechanisms. Such decomposition would be a downstream biological analysis rather than part of the definition or validation of C.

Studies establishing that malignant glioma states shift in relative proportion under perturbation, injury, or spatial context (Couturier et al., 2020; Castellan et al., 2021; Hara et al., 2021; Richards et al., 2021; Ravi et al., 2022; Wang et al., 2022; Doroszko et al., 2025) ask a compositional question: which states expand or contract. C and K were designed to separate that question from a directional one: whether the states that remain present move through transcriptional space in the same direction. The weak and inconsistent behavior of K is consistent with state composition being only loosely coupled to treatment in this dataset at the cell numbers available, which does not contradict the compositional shifts reported elsewhere (Richards et al., 2021; Wang et al., 2022; Doroszko et al., 2025) but indicates that a resolved compositional signal was not detected here. Panobinostat, an HDAC inhibitor with documented antiproliferative activity in glioma (Sears et al., 2021), and RSL3, a GPX4 inhibitor that induces ferroptosis in glioblastoma cells (Li et al., 2021), act through unrelated mechanisms and were both among the nine replicated drugs; the central observation here is that response direction was coherent across states in essentially every comparison, and this coherence was largely specific to the drug rather than to a state or to a general perturbation axis.

The cross-drug comparison most directly constrains the interpretation of C. Pair-level within-drug coherence (median 0.652) exceeded between-drug coherence for different states (median 0.233) in all 52 within-unit drug pairs. More importantly, the result persisted after accounting for the clustering of those pairs: the within-minus-between difference was positive in all 11 TissueIDs, with a median unit-level difference of 0.291 (95% bootstrap CI 0.201-0.495; one-sided exact sign-test P=0.000488). The complementary same-state comparison was also positive overall, although less uniformly, with 9 of 11 TissueIDs positive and a median difference of 0.196 (95% CI 0.057-0.430; P=0.0327). These results argue against the observed C values being explained entirely by a single shared stress or dissociation axis identical across perturbations. As a direct check, a per-cell dissociation-stress score (van den Brink et al., 2017) was computed from 11 canonical stress-response genes and correlated with C across all 43 comparisons: the correlation was weak and not significant (r=-0.183, P=0.239), with no indication that higher stress-score comparisons produced higher C. A shared component of smaller magnitude remains compatible with the positive between-drug similarities and cannot be excluded by this analysis.

There are several limits to what can be concluded from this dataset. Treated and vehicle cells come from matched or adjacent ex vivo slices rather than from the same cells followed through time, and state assignment is recalculated after treatment. A centroid shift can therefore contain both transcriptional movement within a program and redistribution of cells across state boundaries; it is not a lineage trajectory or transition rate. TissueID was used because it reconciles the loom with the 19 reported biological units, but the released metadata do not provide a second patient key with which to verify one-to-one identity. Individual drugs are represented by only three to eight units, and some TissueIDs contribute more than one drug condition. For that reason, drug-level intervals and cross-analysis correlations are descriptive summaries rather than a basis for fine drug ranking. The released metadata provide only cell type, sequencing name, TissueID and treatment label; no dose, concentration or exposure-time field is available, so a dose or exposure-time confound cannot be assessed or excluded from this dataset. A mechanism-of-action stratification of the cross-drug comparison was attempted but is not reported quantitatively: the panel of nine drugs yields only three same-mechanism pairs, all one drug combination (etoposide and topotecan), which is too few to distinguish a mechanism-specific effect from sampling variation. The 500-gene analysis changes the expression coordinates but remains an internal feature-space sensitivity analysis, and the high-confidence, Neftel-module and cross-fit analyses reduce dependence on one hard labeling scheme without converting the states into continuous probabilities. A continuous or probabilistic formulation would therefore test an additional aspect of state representation rather than correct a demonstrated failure of the present result.

The strongest claim supported here is consequently comparative: relative response-direction coherence is more stable across alternative state representations than apparent contraction or expansion of the state configuration. Cross-fitting addresses gene-set overlap, but it reuses the same cells and is not external replication. The vehicle-split and label-permutation nulls indicate that observed C exceeds sampling variation in centroid estimation, while the cross-drug analysis shows that the drug-specific coherence contrast remains positive after reducing the nested 52 drug-pair comparisons to 11 TissueID-level summaries. Neither analysis, however, substitutes for an experimental replicate-vehicle design, so technical slice-to-slice variation independent of drug identity is not fully isolated from the drug-specific effect. Nor is there a functional readout of viability, clonogenicity, recurrence or clinical response. C is therefore a transcriptional response phenotype, not an efficacy score. Independent perturbation data with replicate vehicle slices would test transferability across experiments rather than repair a missing claim in the present analysis. A future extension could also replace hard argmax labels with a continuous or probabilistic state representation and ask whether the same directional relationship is preserved. In practice, C offers a way to triage or group compounds by the consistency of their multi-state transcriptional response — for example, flagging drugs whose effect is concentrated in one state from those that act coherently across the malignant-state landscape — ahead of functional follow-up, rather than a measure of therapeutic efficacy itself. A mechanism-of-action stratification was attempted (Methods) but could not distinguish a mechanism-specific signal from a generic perturbation axis in this dataset, since the drug panel yields only three same-mechanism pairs, all one drug combination (etoposide and topotecan); a drug panel with broader mechanistic replication would be needed to test this question with adequate power.

## Conclusion

A simple geometric framework is introduced to compare drug responses across recurrent malignant glioma states by separating response direction (C), change in state separation (K), and response magnitude (M). Across 43 matched treatment-vehicle comparisons, C was consistently positive, exceeded chance expectation under two within-unit null models, and remained comparatively stable across alternative state representations. In the multi-drug units, within-drug coherence exceeded between-drug coherence in all 11 TissueIDs after pairwise contrasts were collapsed to one summary per biological unit. K was less reproducible when the feature space changed and is therefore interpretable only within an explicitly stated representation. The main contribution is not identification of an effective drug, but a way to quantify whether different glioma states respond coherently to the same perturbation, and a demonstration that this coherence can be tested rather than assumed.

## Ethics statement

This study is a secondary computational analysis of publicly released single-cell data. No participants were recruited and no tissue was collected or experimentally manipulated for this reanalysis. Ethical approval and informed-consent procedures for the original tissue collection are described in the source studies (Zhao et al., 2021; Levitin et al., 2023).

## Data availability

The drug-perturbed glioma single-cell data were generated in the source studies (Zhao et al., 2021; Levitin et al., 2023). GSE148842 is available from the NCBI Gene Expression Omnibus, and the integrated loom analyzed here is publicly linked from the consensus scHPF study (Levitin et al., 2023).

## Code availability

The MATLAB analysis codes are available at: https://github.com/bigbio-el/Glioma-drug-response-geometry.

## Author contributions

Evgeniia Lavrenteva: Conceptualization, Methodology, Software, Formal analysis, Writing – original draft. Kim Kyung Do: Supervision, Writing – review & editing.

## AI declaration

AI-assisted tools were used for language refinement. The authors reviewed and approved the final manuscript.

## Acknowledgements

The authors thank the investigators who generated and publicly released the glioma slice-culture single-cell datasets.

