## Supplementary Materials for "Single-cell geometry of drug responses across malignant glioma states"

### Supplementary information

**Supplementary Figure S1.** Comparison of primary and 500-HVG estimates of K. Primary estimates of change in state separation are compared with estimates obtained in the independently selected 500-gene feature space.

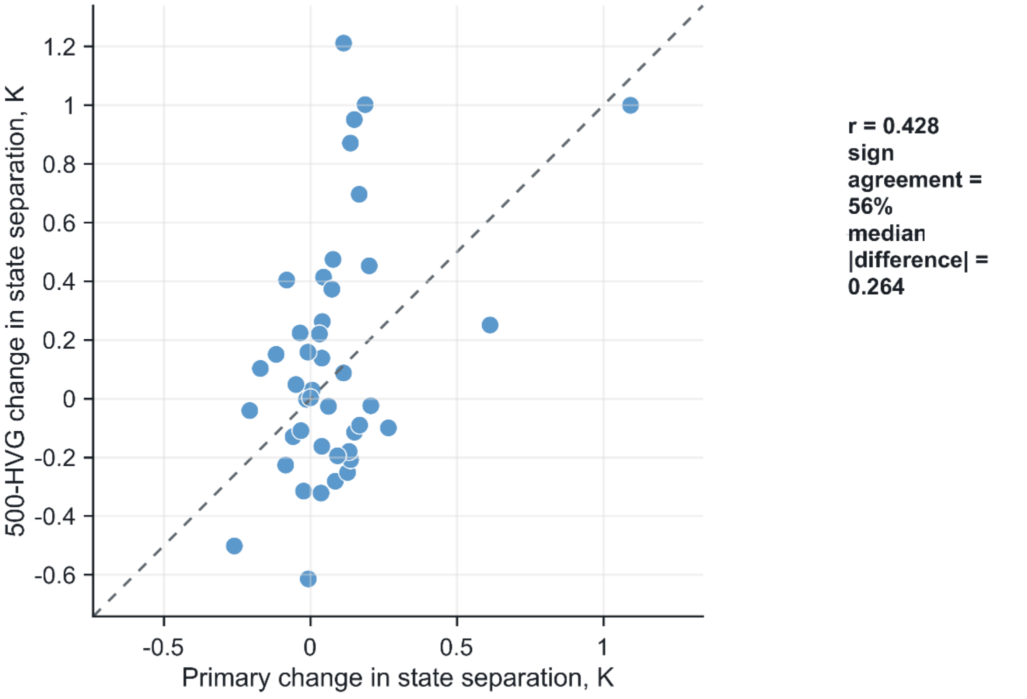


**Supplementary Figure S2.** C and K across all treatment-vehicle comparisons. Response-direction coherence and change in state separation are shown for all 43 biological-unit-drug comparisons. Replicated-drug medians are indicated separately.

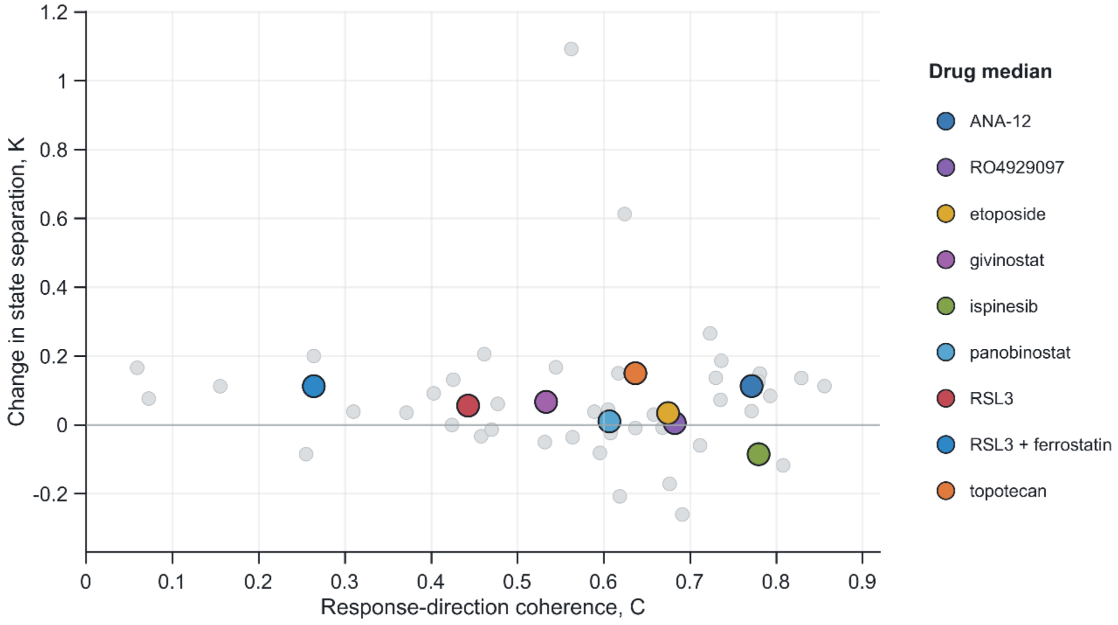

**Supplementary Figure S3.** Agreement between compact and full Neftel state assignments. Agreement of hard state assignments obtained with the compact marker panel and the published Neftel modules.

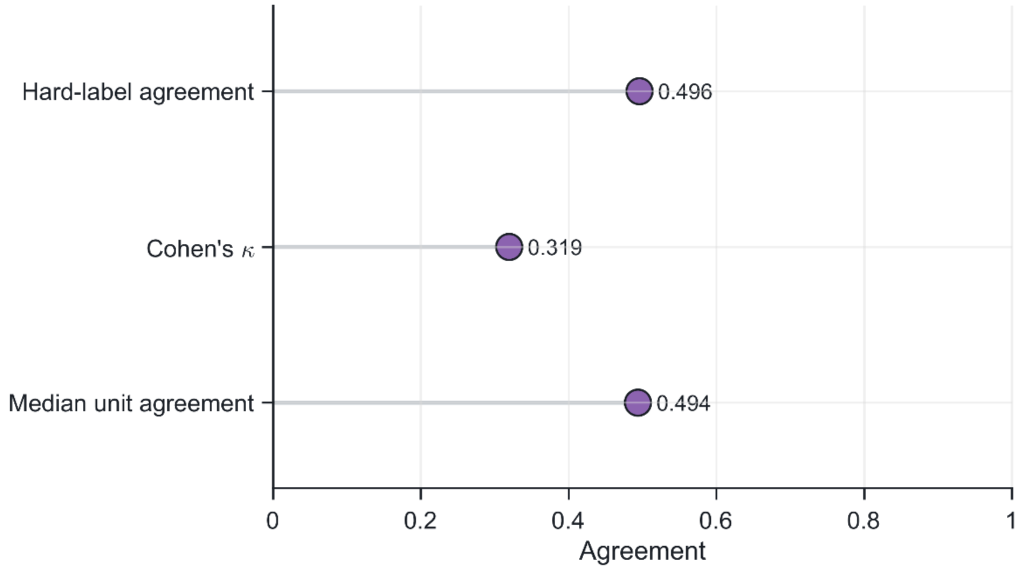

**Supplementary Table S1.** Response-geometry estimates for all biological-unit-drug comparisons. Values of C, K, M and state-composition shift for the 43 matched treatment-vehicle comparisons.

| TissueID | Drug | Vehicle cells | Treated cells | States retained | States used | C | K | M | State shift (TV) |
|---|---|---|---|---|---|---|---|---|---|
| TB5884 | ANA-12 | 10444 | 2045 | 4 | AC,OPC,NPC,MES | 0.856 | 0.113 | 1.727 | 0.169 |
| TB5884 | RO4929097 | 10444 | 2165 | 4 | AC,OPC,NPC,MES | 0.829 | 0.136 | 1.284 | 0.105 |
| TB5884 | etoposide | 10444 | 1631 | 4 | AC,OPC,NPC,MES | 0.736 | 0.186 | 1.482 | 0.110 |
| TB5884 | ispinesib | 10444 | 841 | 4 | AC,OPC,NPC,MES | 0.781 | 0.149 | 1.603 | 0.158 |
| TB5884 | panobinostat | 10444 | 4609 | 4 | AC,OPC,NPC,MES | 0.624 | 0.613 | 2.583 | 0.119 |
| TB5886 | etoposide | 1203 | 2663 | 4 | AC,OPC,NPC,MES | 0.793 | 0.084 | 1.508 | 0.249 |
| TB5886 | ispinesib | 1203 | 3013 | 4 | AC,OPC,NPC,MES | 0.779 | 0.126 | 1.175 | 0.163 |
| TB5944 | ANA-12 | 3541 | 3204 | 4 | AC,OPC,NPC,MES | 0.730 | 0.137 | 1.211 | 0.143 |
| TB5944 | etoposide | 3541 | 1509 | 4 | AC,OPC,NPC,MES | 0.310 | 0.039 | 0.793 | 0.060 |
| TB5944 | ispinesib | 3541 | 857 | 4 | AC,OPC,NPC,MES | 0.255 | -0.085 | 0.848 | 0.115 |
| TB5944 | panobinostat | 3541 | 925 | 4 | AC,OPC,NPC,MES | 0.425 | 0.132 | 1.637 | 0.144 |
| TB5944 | topotecan | 3541 | 1042 | 4 | AC,OPC,NPC,MES | 0.668 | -0.008 | 1.706 | 0.233 |
| TB5966 | RO4929097 | 12968 | 7271 | 4 | AC,OPC,NPC,MES | 0.682 | 0.005 | 0.940 | 0.047 |
| TB5966 | etoposide | 12968 | 13803 | 4 | AC,OPC,NPC,MES | 0.691 | -0.260 | 1.096 | 0.118 |
| TB5966 | ispinesib | 12968 | 11321 | 4 | AC,OPC,NPC,MES | 0.618 | -0.207 | 0.879 | 0.040 |
| TB5966 | panobinostat | 12968 | 1802 | 4 | AC,OPC,NPC,MES | 0.676 | -0.171 | 1.333 | 0.083 |
| TB5966 | topotecan | 12968 | 6315 | 4 | AC,OPC,NPC,MES | 0.617 | 0.150 | 2.265 | 0.270 |
| TB5974 | ANA-12 | 7876 | 2842 | 4 | AC,OPC,NPC,MES | 0.771 | 0.040 | 1.045 | 0.157 |
| TB5974 | RO4929097 | 7876 | 2455 | 4 | AC,OPC,NPC,MES | 0.595 | -0.081 | 1.258 | 0.239 |
| TB5974 | etoposide | 7876 | 3567 | 4 | AC,OPC,NPC,MES | 0.658 | 0.030 | 0.944 | 0.079 |
| TB5974 | ispinesib | 7876 | 9350 | 4 | AC,OPC,NPC,MES | 0.808 | -0.117 | 1.358 | 0.244 |

| | | | | | | | | | |
|---|---|---|---|---|---|---|---|---|---|
| TB5974 | panobinostat | 7876 | 1993 | 4 | AC,OPC,NPC,MES | 0.605 | 0.044 | 1.107 | 0.126 |
| TB5974 | topotecan | 7876 | 4541 | 4 | AC,OPC,NPC,MES | 0.636 | -0.009 | 0.828 | 0.067 |
| TB5980 | topotecan | 2180 | 3614 | 4 | AC,OPC,NPC,MES | 0.723 | 0.266 | 1.385 | 0.179 |
| TB6140 | panobinostat | 5306 | 821 | 4 | AC,OPC,NPC,MES | 0.531 | -0.050 | 1.137 | 0.223 |
| TB6181 | rsl3 | 5426 | 1461 | 4 | AC,OPC,NPC,MES | 0.461 | 0.206 | 0.989 | 0.222 |
| TB6181 | rsl3_ferrostatin | 5426 | 718 | 4 | AC,OPC,NPC,MES | 0.264 | 0.200 | 0.903 | 0.137 |
| TB6224 | givinostat | 6752 | 1379 | 4 | AC,OPC,NPC,MES | 0.477 | 0.061 | 0.660 | 0.097 |
| TB6328 | rsl3 | 3501 | 4317 | 4 | AC,OPC,NPC,MES | 0.711 | -0.060 | 0.901 | 0.142 |
| TB6328 | rsl3_ferrostatin | 3501 | 5122 | 4 | AC,OPC,NPC,MES | 0.458 | -0.033 | 0.612 | 0.128 |
| TB6393 | etoposide | 1652 | 865 | 4 | AC,OPC,NPC,MES | 0.470 | -0.013 | 0.831 | 0.073 |
| TB6393 | panobinostat | 1652 | 1687 | 4 | AC,OPC,NPC,MES | 0.608 | -0.024 | 0.930 | 0.045 |
| TB6458 | rsl3 | 2901 | 6381 | 4 | AC,OPC,NPC,MES | 0.371 | 0.036 | 0.870 | 0.129 |
| TB6458 | rsl3_ferrostatin | 2901 | 1013 | 4 | AC,OPC,NPC,MES | 0.155 | 0.113 | 0.696 | 0.085 |
| TB6488 | givinostat | 3647 | 181 | 3 | AC,OPC,MES | 0.059 | 0.166 | 1.065 | 0.097 |
| TB6505 | givinostat | 28661 | 4494 | 4 | AC,OPC,NPC,MES | 0.735 | 0.073 | 0.740 | 0.036 |
| TB6528 | rsl3 | 1564 | 1158 | 4 | AC,OPC,NPC,MES | 0.403 | 0.092 | 1.102 | 0.133 |
| TB6534 | rsl3 | 1491 | 771 | 4 | AC,OPC,NPC,MES | 0.424 | 0.000 | 1.001 | 0.088 |
| TB6545 | givinostat | 17741 | 2832 | 4 | AC,OPC,NPC,MES | 0.589 | 0.038 | 0.837 | 0.096 |
| TB6545 | rsl3 | 17741 | 498 | 3 | AC,OPC,NPC | 0.562 | 1.092 | 3.648 | 0.408 |
| TB6813 | rsl3 | 2722 | 610 | 4 | AC,OPC,NPC,MES | 0.072 | 0.077 | 0.733 | 0.055 |
| TB6814 | rsl3 | 1393 | 652 | 4 | AC,OPC,NPC,MES | 0.564 | -0.036 | 0.775 | 0.138 |
| TB6814 | topotecan | 1393 | 2799 | 4 | AC,OPC,NPC,MES | 0.544 | 0.168 | 1.111 | 0.136 |

**Supplementary Table S2.** Genes used for compact malignant-state assignment. The 34-gene marker panel used to score AC-like, OPC-like, NPC-like and MES-like states.

| State | Genes |
|---|---|
| AC-like | CST3, GFAP, S100B, HOPX, SLC1A3, MLC1, AQP4 |
| OPC-like | PLP1, ALCAM, OLIG1, OLIG2, OMG, PLLP, PDGFRA |
| NPC-like | SOX4, SOX11, DCX, CD24, DLL3, STMN2, STMN4, RND3, DLX5, DLX6-AS1 |
| MES-like | VIM, ANXA1, ANXA2, CHI3L1, CD44, DDIT3, ADM, HILPDA, ENO2, LDHA |

**Supplementary Table S3.** Comparison of results across sensitivity analyses. Descriptive concordance of C and K after high-confidence state filtering, analysis in the independent 500-HVG space, use of the full Neftel modules and non-overlapping gene cross-fitting.

| Analysis | Metric | n pairs | Pearson r | Sign agreement | Median abs. change | Median signed change | Median abs. difference | Median signed difference |
|---|---|---|---|---|---|---|---|---|
| High-confidence compact states | C | 42 | 0.931 | 1.000 | 0.168 | -0.168 | - | - |
| High-confidence compact states | K | 42 | 0.918 | 0.881 | 0.048 | 0.023 | - | - |
| Independent 500-HVG geometry | C | 43 | 0.634 | 1.000 | - | - | 0.258 | 0.258 |
| Independent 500-HVG geometry | K | 43 | 0.428 | 0.558 | - | - | 0.264 | 0.003 |
| Full published Neftel modules | C | 43 | 0.783 | 1.000 | - | - | 0.103 | 0.087 |
| Full published Neftel modules | K | 43 | 0.639 | 0.698 | - | - | 0.125 | 0.046 |
| Neftel non-overlapping cross-fit | C | 43 | 0.809 | 1.000 | - | - | 0.148 | 0.148 |
| Neftel non-overlapping cross-fit | K | 43 | 0.546 | 0.628 | - | - | 0.211 | 0.004 |
